# Brain coding of social network structure

**DOI:** 10.1101/850065

**Authors:** Michael Peer, Mordechai Hayman, Bar Tamir, Shahar Arzy

## Abstract

To successfully navigate our social world, we keep track of other individuals’ relations to ourselves and to each other. But how does the brain encode this information? To answer this question, we mined participants’ social media (Facebook^TM^) profiles to objectively characterize the relations between individuals in their real-life social networks. Under fMRI, participants answered questions on each of these individuals. Using representational similarity analysis, we identified social network structure coding in the default-mode network (medial prefrontal, medial parietal and lateral parietal cortices). When regressing out subjective factors (ratings of personal affiliation, appearance and personality), social network structure information was uniquely found in the retrosplenial complex, a region implicated in spatial processing. In contrast, information on individuals’ personality traits and affiliation to the subjects was found in the medial prefrontal and parietal cortices, respectively. These findings demonstrate a cortical division between representation of structural, trait-based and self-referenced social knowledge.

## Introduction

Social interactions and connections form a major part of human life (Dunbar, 2018). In order to successfully operate in a social world, humans need to represent information about hundreds of individuals comprising their social network – how these individuals are affiliated to themselves, what are their relations with each other, their membership in different social groups, and subjective evaluations of features such as their personality traits. While the brain systems related to personal affiliation and trait judgments have been investigated (Hassabis et al., 2014; Jenkins and Mitchell, 2011; Jenkins et al., 2008; Maddock et al., 2001; Mitchell et al., 2006; Parkinson et al., 2014, 2017; Shah et al., 2001; Wlodarski and Dunbar, 2016), it is less clear how humans store information on the mutual relations between other people in their social network. Furthermore, as the multiple aspects comprising social knowledge may be correlated with each other, it is unclear how the brain simultaneously represents these different kinds of information. Finally, it has been suggested that social networks and spatial environments may share similar representations (Epstein et al., 2017; Parkinson and Wheatley, 2013; Peer et al., 2015), but evidence for social information processing in spatial regions is lacking.

Several neuroimaging lines of research may shed light on how the brain codes different features of known others, including their relation to the self. Investigations of the brain coding of personal affiliation mostly compared thinking of familiar others at different levels of personal proximity (increased familiarity / closer relationship), demonstrating that differences pertaining to personal affiliation level mostly occur in the medial and lateral parietal and temporal regions (Maddock et al., 2001; Parkinson et al., 2014, 2017; Shah et al., 2001; Tavares et al., 2015; Wlodarski and Dunbar, 2016). Other studies have looked at neural correlates of overall social network size using social media information, showing that network size is correlated with the size and connectivity of medial temporal and orbitofrontal structures (Bickart et al., 2012; Hampton et al., 2016; Von Der Heide et al., 2014; Kanai et al., 2011; Powell et al., 2012). Investigations of how the brain codes others’ personality traits implicated the medial prefrontal cortex in trait judgments and trait differences (Hassabis et al., 2014; Jenkins and Mitchell, 2011; Jenkins et al., 2008; Mitchell et al., 2006). These findings converge on regions comprising the brain’s default-mode network, known to be involved in self-referential processing and internal mentation (Buckner and DiNicola, 2019; Buckner et al., 2008), as well as in social mentation about others (e.g. Theory of Mind (Buckner and Carroll, 2007; Saxe and Kanwisher, 2003)).

More recent studies have directly investigated aspects of the brain coding of social network structure. Studies of how the brain represents social hierarchy and power (using fictional individuals) demonstrated involvement of the hippocampus, amygdala, medial prefrontal and posterior cingulate cortices (Kumaran et al., 2012, 2016; Tavares et al., 2015). Another recent study mapped real-life relations between people in a specific class of students, identifying coding of personal affiliation to the subject and measures of individuals’ centrality in the social network (eigenvector centrality and brokerage) across regions of the lateral temporal, medial and lateral parietal and medial prefrontal cortex (Parkinson et al., 2017). These studies provide important insights regarding the brain systems involved in representation of social relations. However, several questions remain unanswered: 1) how does the brain represent the actual allocentric structure of each person’s social network (i.e. the mutual relations between other individuals)? 2) Is this representation independent of other factors, such as egocentric affiliation to the self, or similarity between people along different features? 3) How does the brain represent the structure of real-world, large-scale social networks, that may contain individuals from different groups and social contexts?

To tackle these issues, we extracted mutual friendship information from subjects’ social media profiles. We used this information to calculate the proximity between all individuals in each subject’s network (by their proportion of mutual friends relative to all friends) and reconstruct the network’s structure, independently of these individuals’ affiliation to the subject. We then recorded functional MRI activity while subjects thought about each individual. To encourage thinking about different individuals, in each trial the name of one person was presented, and the subject was asked to answer a question about this person by providing a rating (regarding personality traits, appearance, or personal affiliation to themselves; Figure 1). While these questions served mainly to encourage mentalizing about each individual, they also provided subjective ratings, that enabled calculating the similarity between these individuals in terms of their subjectively perceived characteristics. Importantly, subjects were not explicitly asked at any stage about relations between individuals along any social dimension. Finally, we used representational similarity analysis (RSA) of multi-voxel activity patterns to identify brain regions representing unique information on different aspects of social knowledge – social proximity in the network, personal affiliation, personality and appearance. The use of RSA relies on the assumption that if a brain region codes information on a specific dimension (here: social network proximity), it will display similar activity patterns when subjects think of individuals who are close along this dimension. By concurrently investigating the similarity between individuals along multiple social dimensions, our study enables dissociating the unique aspects comprising relational social knowledge.

**Figure 1:**
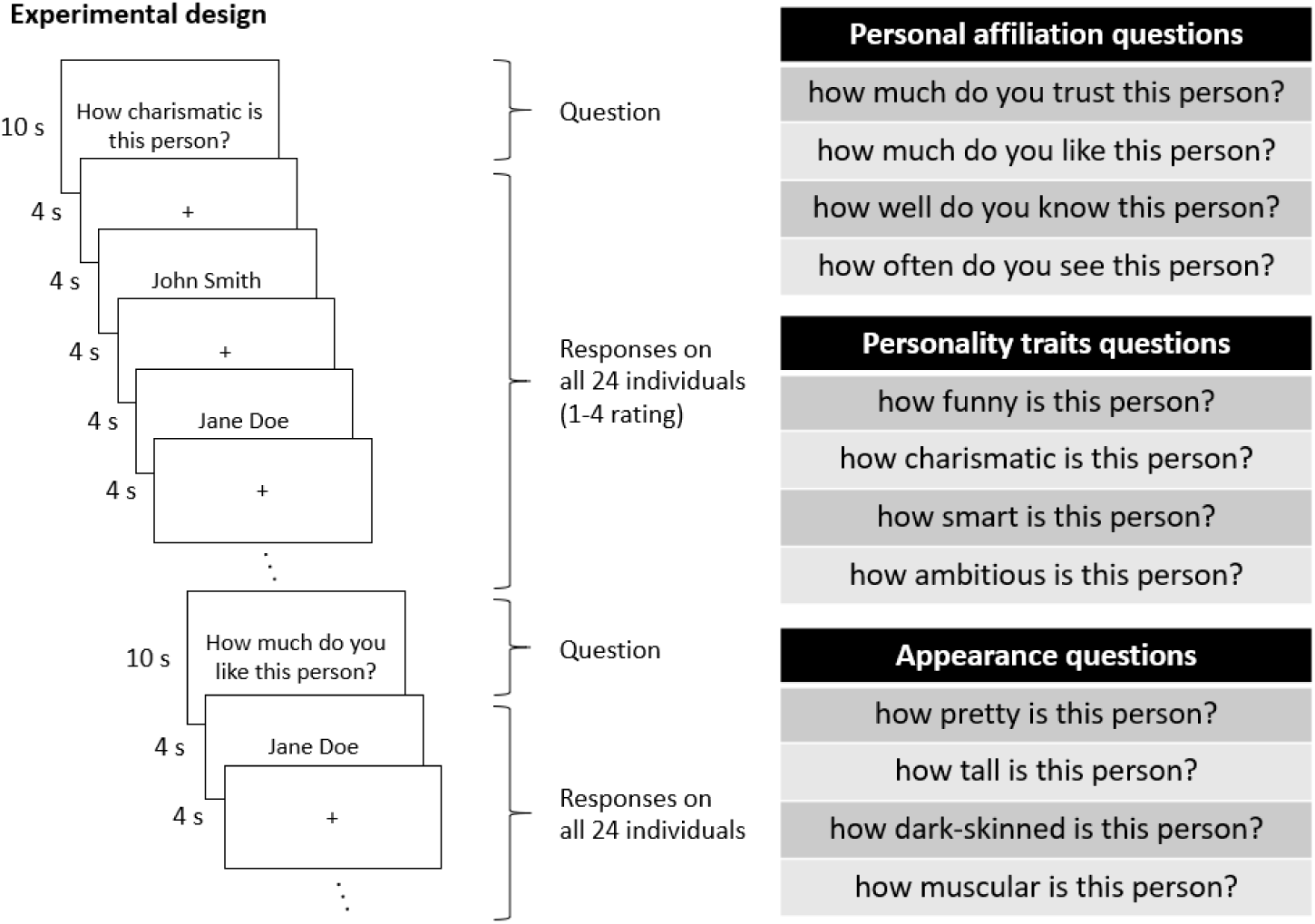
Experimental design. The paradigm was designed to make subjects mentalize about people they know. To this end, subjects saw a question, followed by the names of 24 individuals from their personal social network, and were asked to respond to the same question on each of the individuals using a rating of 1-4. The questions were designed to encourage mentalizing about each individual in turn. Each of the twelve questions was presented for ten seconds, followed by a randomly ordered sequence of all 24 names, each presented for four seconds with an interval of four seconds between names. The same 24 unique individual names were used for all questions.

## Results

### The default mode network represents information on social network distance between personally-familiar people

Social network distances between individuals were computed using their objective friendship patterns as extracted from social media, independent of their affiliation to the subjects. Representational similarity searchlight analysis was then used to compare social network distances to the similarity in neural patterns when subjects think of these individuals. This analysis revealed coding of social network distances in regions within the medial and lateral parietal and frontal cortices (Fig. 2). Comparison of the regions coding social network distances to large-scale resting-state networks (Yeo et al., 2011) reveals that these regions correspond to the default-mode network (Dice’s coefficient for default-mode network overlap = 0.79; p<0.001, permutation test; no other resting-state network had significant overlap with social network distance coding regions). To verify that the results do not depend on the specific social distance measure used (proportion of mutual friends), the analysis was repeated using other social network distance measures - shortest path length, total number of mutual friends, direct connectivity and communicability (see Figure S1 legend). All measures except for communicability yielded similar results (Figure S1), demonstrating that the coding of social network distances in the default-mode network is robust across distance measures.

**Figure 2:**
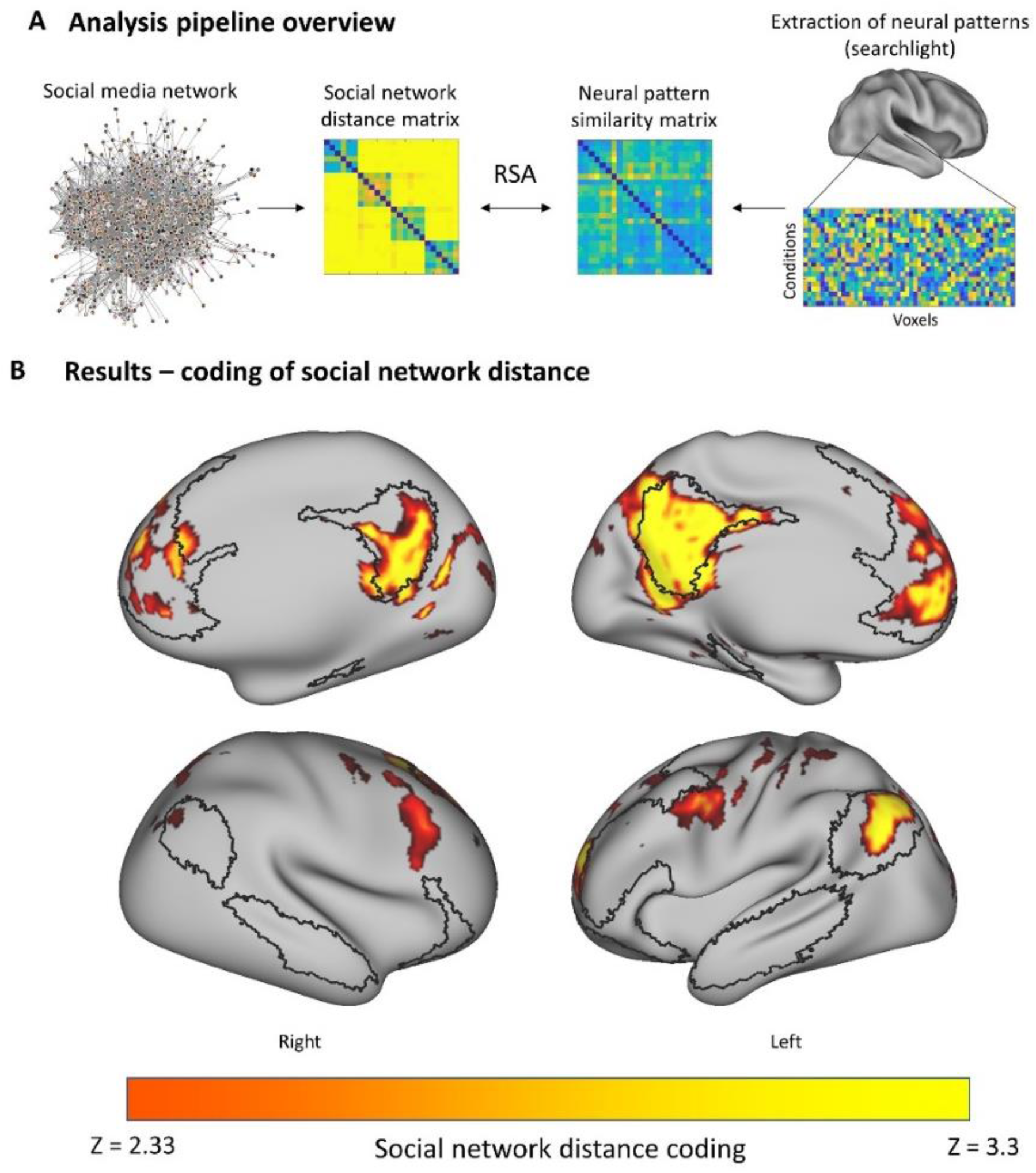
the default-mode network represents social network structure information. Results of representational similarity searchlight on social media network proximity between individuals. Pattern similarity across regions of the default-mode network was associated with proximity between individuals in each subject’s personal social network, as derived from the social media connectivity data. The black outline delineates the default-mode network, as defined in Yeo et al., 2011. All p-values<0.01, Monte-Carlo permutation test, threshold-free cluster enhancement correction for multiple comparisons.

### Dissociation between coding of social network structure, personal affiliation, and personality traits in the parietal and frontal cortices

The correspondence between neural pattern similarity and social network distance could be related to other factors that are correlated with social network distance, such as personal affiliation to the subject and similarity in personality traits (Table S1). To dissociate these factors, we constructed similarity matrices for individuals’ personality traits, appearance, and personal affiliation to the subject, based on subjects’ subjective ratings. Representational similarity searchlight using these matrices revealed that personal affiliation and personality similarity (but not appearance) correspond to neural pattern similarity within the default-mode network, in accordance with their shared variance with social network proximity (Table S1, Figure S2). To measure the independent contribution of each factor (similarity in activity explained by the unique variance of each factor, excluding the effect of the common variance), we regressed these matrices from one another and performed an RSA searchlight on the residuals (Figures 3, S3). We found that after regression of the abovementioned factors, the only region representing information on social network structure is the posterior part of the medial parietal lobe. An adjacent and partially overlapping region within the medial parietal lobe, and a region in the lateral parietal lobe, represented information on personal affiliation to the subject, while information on similarity in individuals’ personality traits was found only in the medial prefrontal lobe (Figure 3). No region represented information on similarity in personal appearance.

**Figure 3:**
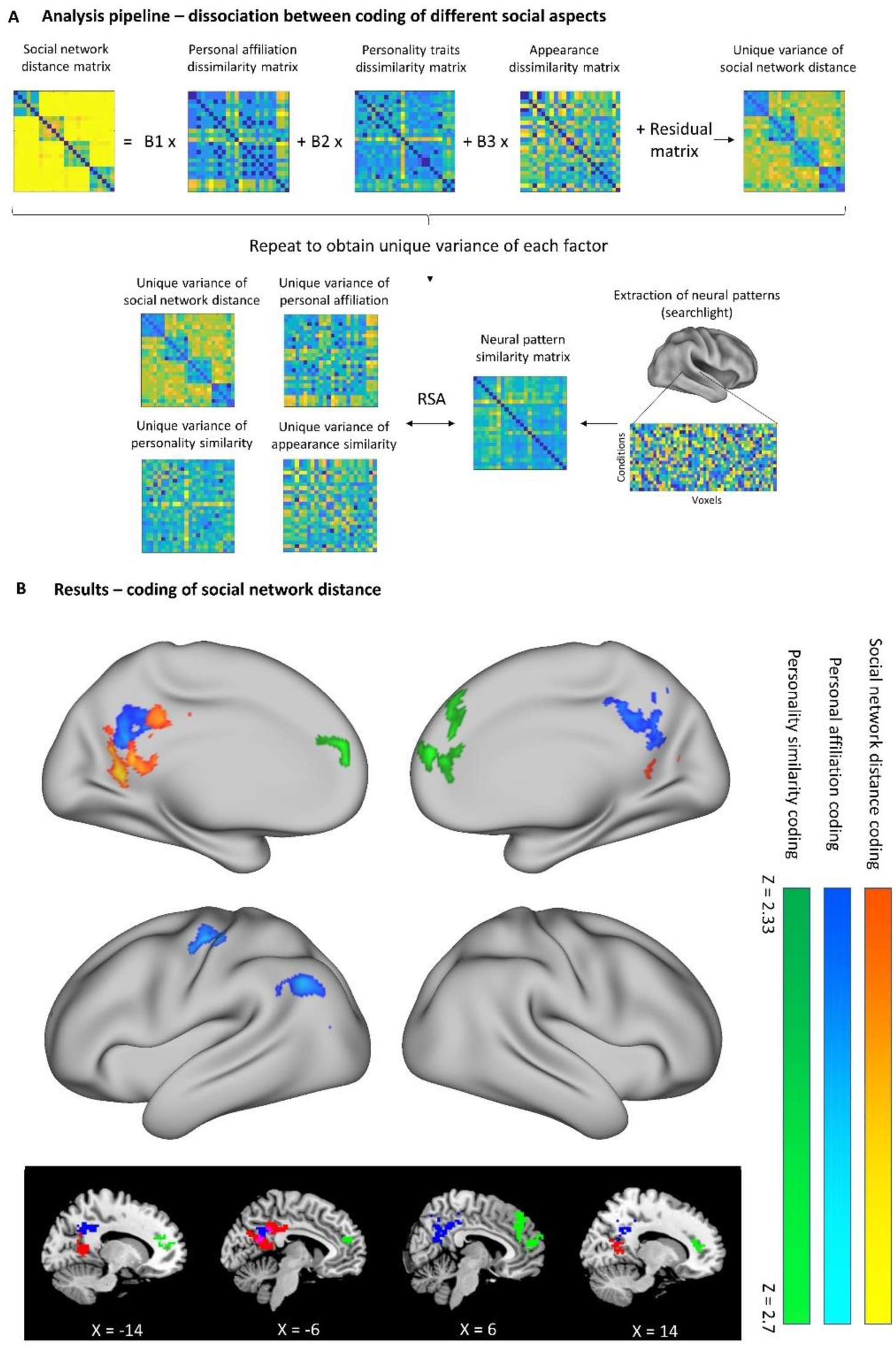
Dissociation between representations of social network structure, personal affiliation and personality traits. A) Similarity matrices for individuals’ social network distance, personal affiliation to the subject, personality traits, and appearance were regressed from one another, and the residual matrix was used in a representational similarity searchlight, to identify the independent variance explained by each factor. B) Along the medial wall, activity pattern similarity in the medial parietal lobe represented information on the social proximity of individuals to one another and their personal affiliation to the subject, while pattern similarity in the medial prefrontal cortex was correlated to similarity between individuals in their perceived personality traits. No voxels were found significantly containing information on similarity in appearance. Representational similarity searchlight, spherical radius = 3 voxels, p<0.01, Monte-Carlo permutation test, threshold-free cluster enhancement correction for multiple comparisons.

### Social network structure information is found in spatial processing regions

The identified medial parietal region coding for social network structure appears to overlap with the retrosplenial complex (RSC), a region implicated in spatial mapping (Epstein, 2008; Epstein et al., 2007). The social network coding region also appears to overlap with regions we identified in a previous study (Peer et al., 2015) that are active when subjects make egocentric proximity comparisons to different places (“spatial orientation region”) and to different people (“social orientation region”). To measure whether these regions indeed represent social network information, we performed the RSA analysis in each of these independently defined regions of interest, for each social factor of interest (social network proximity, personal affiliation, personality and appearance) after regressing out the other factors. We found significant coding of social network distance between individuals in the left and right retrosplenial complex, as well as within the spatial and social orientation regions (representational similarity analysis, all p-values<0.05, FDR-corrected for multiple comparisons across regions of interest; Figure 4). Coding of personal affiliation to the subject was also found in the social orientation region, but not in the other regions of interest (representational similarity analysis, all p-values<0.05, FDR-corrected for multiple comparisons across regions of interest; Figure 4). No significant effects were found in other spatial processing brain regions (parahippocampal place area, occipital place area, or hippocampus), and no coding was found in any region of interest for similarity in personality or appearance. These findings indicate that the retrosplenial complex, as identified by an independent localizer for spatial scene selectivity (Julian et al., 2012), contains information on social network relations between individuals.

**Figure 4:**
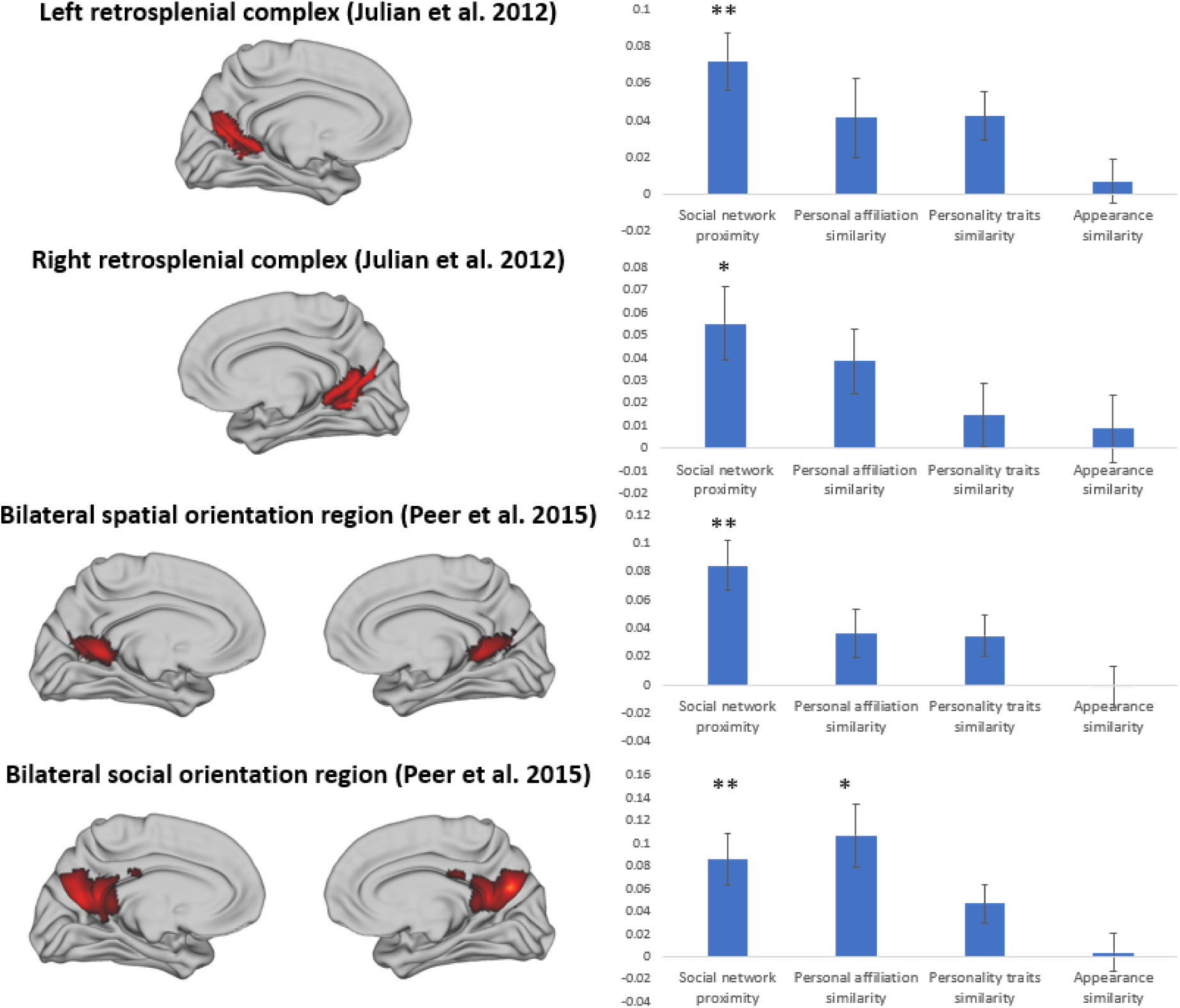
Social network structure is coded in spatial processing regions. Representational similarity analysis was performed in regions of interest defined using scene viewing (retrosplenial complex; Julian et al., 2012) and spatial or social self-proximity judgments (spatial and social orientation regions; Peer et al., 2015). The analysis of the coding of each factor (social network proximity, personal affiliation, personality and appearance) was performed after regression of the matrices corresponding to the other factors, as in Figure 3. Significant coding of social network proximity between individuals was found in the right and left RSC, and in the spatial and social orientation regions. Significant coding of personal affiliation was also found in the social orientation region. (* - p<0.05, ** - p<0.01, all p-values FDR-corrected for multiple comparisons across regions).

## Discussion

Using objective information from subjects’ social media profiles and subjects’ subjective ratings of individuals’ characteristics, we dissociated regions representing different aspects of social knowledge. Activity pattern similarity within the default-mode network predicted social network proximity between individuals that the subjects mentalized about. Activity patterns in the default-mode network also reflected individuals’ personal affiliation to the subject as well as their personality traits. When dissociating these dimensions of social similarity, information on social network structure was identified only in the medial parietal lobe. In contrast, information on the personal affiliation of individuals to the subject was represented in the medial and lateral parietal cortex, and information on individuals’ personality traits was found in the medial prefrontal cortex. Notably, representation of the relations between individuals across the social network was evoked spontaneously in the recorded brain activity, despite subjects not being explicitly asked about these relations during the experiment. Finally, the region coding for relations across the social network overlapped with the retrosplenial complex, a spatial processing region, suggesting that this region might have a general role in mapping relations across the spatial and social cognitive domains.

Our findings suggest that knowledge about familiar others can be dissociated into at least three components, that are preferentially processed in different cortical circuits – knowledge of others’ personality traits (medial prefrontal cortex), knowledge of affiliation of others to the self (medial and lateral parietal cortex), and knowledge of the higher-order organization of the social network (medial parietal - retrosplenial cortex). All of these regions are part of the default-mode network, that is related to the processing of social and self-referential information (Buckner et al., 2008). Previous studies using univariate and multivariate methods observed personal-affiliation related activity in the medial and lateral parietal cortex (Gusnard et al., 2001; Maddock et al., 2001; Parkinson et al., 2014, 2017; Peer et al., 2015; Shah et al., 2001; Tavares et al., 2015; Wlodarski and Dunbar, 2016), as well as personality-trait judgment related activity in the medial prefrontal cortex (Chavez and Heatherton, 2015; Hassabis et al., 2014; Mitchell et al., 2006). Our findings extend these findings to show coding of real-life social distances between familiar people in the medial parietal cortex, in partial dissociation from other types of social knowledge. Overall, the default-mode network appears to have an overarching role in processing information about familiar others, with sub-specializations for different types of information in its different subregions.

We observed coding of social network relations at the retrosplenial complex (RSC), a region related to spatial scene processing. The RSC is known to integrate egocentric and allocentric spatial information (Arzy and Schacter, 2019; Byrne et al., 2007). In accordance with this idea, we also observed here coding of ‘egocentric’ social information (self-referenced personal affiliation) and ‘allocentric’ social information (objective relations between others in the social network, irrespective of relation to the self) in the RSC. In a previous study, we identified partially overlapping regions in the medial parietal-RSC region that were active for self-referential judgments in the spatial, social and temporal domains (Peer et al., 2015). A similar convergence of social and spatial information has also been shown in the lateral parietal cortex (Parkinson et al., 2014) and in the medial temporal lobe (Tavares et al., 2015). Additional studies of the medial parietal lobe have demonstrated that it represents relations between social group categories (Leshinskaya et al., 2017) and across events in time (Baldassano et al., 2017). Our findings here strengthen the idea that the brain’s spatial processing system has a domain-general role, and is involved in mapping knowledge across space, time and the social domain (Arzy and Schacter, 2019; Behrens et al., 2018; Bellmund et al., 2018; Constantinescu et al., 2016; Epstein et al., 2017; Parkinson and Wheatley, 2013, 2015; Parkinson et al., 2014; Peer et al., 2015; Schafer and Schiller, 2018; Tavares et al., 2015).

Despite this convergence, our findings somewhat conflict with previous findings on spatial, social and scene processing in the medial parietal lobe. Our previous study identified a division between posterior parietal regions preferentially active for spatial processing and more anterior regions preferentially active for social processing (Peer et al., 2015). This finding was recently replicated and extended by another study that used recall of people and places and found the same posterior-anterior division between spatial and social processing (Silson et al., 2019a). In addition, comparison of scene viewing to other categories identifies the posterior region, retrosplenial complex, as scene-selective (Epstein, 2008; Epstein and Higgins, 2007; Silson et al., 2016, 2019b). The posterior-anterior division is supported by recent resting-state studies that divided the default-mode network into posterior and anterior components (Braga and Buckner, 2017; Braga et al., 2019). In contrast, our findings here suggest that both the posterior and anterior medial parietal cortex represent social information. In the same vein, in a recent study we found that both posterior and anterior medial parietal regions also engage in spatial processing (at different scales) (Peer et al., 2019). These findings suggest that medial parietal subdivisions might not be driven strictly by domain, but by other factors related to domain and task. One possibility is that the posterior region is representing structural relations (e.g. spatial maps and social networks), while the anterior region represents self-affiliation. Other factors that may distinguish posterior and anterior activations may be visual vs. abstract processing and representation, allocentric vs. egocentric processing, or emotionally-significant vs. insignificant relations representation. Furthermore, the relations between posterior and anterior regions may be considered as continuous gradients rather than distinct activations (Buckner and DiNicola, 2019; Huntenburg et al., 2017; Margulies et al., 2016; Peer et al., 2019). Future studies may shed more light on the factors influencing medial parietal lobe selectivity to different types of stimuli in the social, spatial and other cognitive domains.

Our study used information from social media to map real-life, ecologically-valid relations between individuals, which were implicitly invoked by asking subjects to mentalize about each individual separately. Previous neuroscience studies on social relations mostly introduced subjects to new characters with different personality traits (Hassabis et al., 2014; Kumaran et al., 2012; Mitchell et al., 2006; Tavares et al., 2015), or used a small subset of personally familiar names (Jenkins et al., 2008). Past studies using social media information have been mostly limited to studying the brain correlates of social network size (Bickart et al., 2012; Hampton et al., 2016; Kanai et al., 2011). One exception is a recent study which investigated the representation of a real-life social network of students, by mapping all of their inter-relations, and using multivariate methods to investigate characteristics of individuals such as their centrality in the social network and their personal affiliation to the experimental subjects (Parkinson et al., 2017). Here we extended these investigations to provide additional insights into how the brain codes relations between individuals from different social groups and contexts, who are personally known for a prolonged period of time. The combination of information from social media with multivariate pattern analysis is a potentially valuable tool for future investigations of real-life social information representations.

Previous studies demonstrating coding of relational knowledge across cognitive domains highlighted the role of the hippocampus in these processes. The hippocampus represents distances between spatial locations (Deuker et al., 2016), and was also shown to map information about personal affiliation and power (Tavares et al., 2015) and relations between items in other abstract domains (Schapiro et al., 2012). We did not identify coding of social information in the hippocampus in this study, although this null result does not indicate that the information is not there. A possible reason for the null finding is that the hippocampus may primarily represent newly learned relations, in contrast to neocortical regions that may represent information consolidated over time, including life-long social relations, as tested here (Norman and O’Reilly, 2003; Winocur et al., 2007). Another, more technical explanation is that the hippocampus is affected by susceptibility artifacts and signal distortions more than other regions (Olman et al., 2009; Peer et al., 2016), and we did not use here a scanning protocol optimized to image the medial temporal lobe (Weiskopf et al., 2006). Further research is needed to explore the hippocampal role in representation and processing of real-world social network structure.

Our study has several limitations. First, we used information extracted from social media to reconstruct each subject’s personal social network structure. This information may contain inaccuracies with respect to the real-world social structure (e.g. due to people who do not have a social media account, although we verified that this is not the case for the 24 individuals picked for the experiment). Furthermore, we only used data on whether individuals are connected on social media, regardless of the strength or type of this connection. However, these inaccuracies cannot explain the significant coding of social information that we do observe. Third, we asked subjects to provide individuals from several different social groups, and therefore group membership may explain a large amount of variability in the distance matrices, and cannot be completely disambiguated from specific connection between individuals. Fourth, to avoid circularity in the analyses, spatial and social functional regions of interest (e.g. RSC and orientation regions) were defined using masks from separate studies; therefore, these regions may not precisely correspond to each individual’s functional regions. Finally, a shared social context between familiar individuals can be related to other shared contextual factors known to the subject, such as common spatial context or common episodic memories. Further experiments specifically designed to dissociate social and episodic knowledge may provide insights into the unique processes involved in each of these aspects.

In conclusion, we have demonstrated dissociable coding of different aspects of knowledge about personally-familiar others, in different regions of the default mode network, including brain regions implicated in the coding of spatial cognitive maps. Our findings suggest that the brain stores objective information on how other people are related to each other irrespective of the relation of these people to the subject, information that can be useful for appropriate social behavior and interactions. Finally, in combination with previous studies, our findings support the notion that the brain uses the same system to process and store relational knowledge across different cognitive domains, and suggest a division between structural and self-referenced information that may apply in both the spatial and the social domains.

## Methods

### Subjects

Eighteen healthy subjects (nine males, mean age 25.8 ± 3.4 y) participated in the study. Subjects were required to have a Facebook social media account. All subjects provided written informed consent, and the study was approved by the ethical committee of the Hadassah Hebrew University Medical Center.

### Experimental stimuli

Subjects were asked to provide the full names of 24 of their Facebook social media platform friends, from four different social groups in their life (six people from each group). All 24 names had to be distinct from each other. Examples of social groups provided by the subjects are family members, friends from a trip abroad, high school friends, national service friends, friends from work, friends from the university, and friends from the youth movement.

### Extraction of social network structure data

Social network connectivity information was extracted using the Lost Circles Chrome extension (http://lostcircles.com). Lost Circles is a tool that maps which of the subject’s friends on Facebook are connected with each other (“mutual friends”), thus enabling reconstruction of their social network. Each subject installed and ran the Lost Circles extension, and the resulting data file was converted to a binary connectivity matrix between all individuals in the subject’s social network. Subjects’ social network comprised on average of 571±379 individuals (mean±SD, smallest network size = 99, largest network size = 1279). No other data about Facebook interactions between individuals was collected. The experimenters did not have access to subjects’ Facebook accounts at any point.

### Experimental paradigm

During the experiment, subjects were asked to respond to twelve questions about each of the 24 individuals whose names they provided, while undergoing fMRI scanning (Figure 1). Four questions were related to personal affiliation – “how much do you trust this person”, “how much do you like this person”, “how well do you know this person”, “how often do you see this person”. Four questions were related to personality traits – “how funny is this person”, “how charismatic is this person”, “how smart is this person”, “how ambitious is this person”. The last four questions were related to appearance – “how pretty is this person”, “how dark-skinned is this person”, “how tall is this person”, “how muscular is this person”. The questions were designed to encourage deep elaboration of each individual in turn, but in addition provided data (subjects’ subjective ratings) for control analyses of factors influencing brain activity beyond social network structure.

The experiment consisted of six experimental runs, each divided into two question phases (for a total of twelve questions across the experiment). Each question phase started with the presentation of one of the twelve questions for ten seconds, followed by four seconds of fixation. After each question, the 24 names provided by the subject were presented sequentially, each shown for four seconds followed by four seconds of fixation before the next name. Subjects were required to answer the preceding question for all of the 24 names using a four-button response box, leading to a rating of one to four for each response. Question order was randomized across subjects, and the order of the 24 presented names was randomized between question phases.

### Calculation of distance matrices

For each subject, four 24×24 dissimilarity (distance) matrices were computed between each of the 24 subject-provided individual names:

1. Dissimilarity in social network position – the social network distance between each pair of individuals was computed as one minus the proportion of friends they share out of the total number of their friends (Bapna et al., 2017; Granovetter, 1977). Intuitively, individuals with many mutual friends will occupy the same “region” of the social graph (or same social group), while individuals occupying different graph regions will have few mutual friends. This social proximity measure controls for individuals’ popularity (overall number of friends) and diminishes the effects of “shortcuts” between graph regions, as opposed to path length measures. See below for alternative social distance measures.
2. Dissimilarity in personal affiliation to the subject – calculated as the Euclidean distance between the subject’s responses to the four personal affiliation questions. For example, if a subject gave the same answers for two individuals regarding how much she/he trusts them, how frequently they meet, etc., these individuals will have a high similarity value.
3. Dissimilarity in personality – calculated as the Euclidean distance between answers to the four personality traits questions.
4. Dissimilarity in appearance – calculated as the Euclidean distance between answers to the four appearance questions.

In addition to the main Facebook network social distance measure described above (proportion of mutual friends), we also attempted in supplementary analyses to use other measures of social network proximity: the overall number of mutual friends two individuals have, the length of the shortest path length between the individuals (calculated with the Brain Connectivity Toolbox, (Rubinov and Sporns, 2010)), the existence of a direct connection between the individuals (binary measure), and the communicability between individuals as reflected by the matrix exponential (Estrada and Hatano, 2008). All measures were normalized to the range of 0-1 by division by the maximal values and converted to dissimilarity by subtraction from one.

Three subjects had one name each that could not be identified in their Facebook network (one subject provided a name that did not appear in the Facebook friends list, and two subjects had two different individuals in their network with the same first and last name). For these subjects, the missing names were removed from all connectivity matrices and the resulting 23×23 distance matrices were used in all further analysis.

### MRI acquisition

Subjects were scanned in a 3T Siemens Skyra MRI (Siemens, Erlangen, Germany) at the Edmond and Lily Safra Center (ELSC) neuroimaging unit. Blood oxygenation level-dependent (BOLD) contrast was obtained with an echo-planar imaging sequence [repetition time (TR), 2s; echo time (TE), 30ms; flip angle, 75°; field of view, 192mm; matrix size, 64×64; functional voxel size, 3×3×3mm; 37 slices, descending acquisition order, 0.3mm gap; 213 TRs per run]. In addition, T1-weighted anatomical images (1×1×1mm, 160 slices) were acquired for each subject using an MPRAGE protocol [TR, 2,300ms; TE, 2.98ms; flip angle, 9°; field of view, 256mm].

### MRI processing

fMRI data were processed and analyzed using BrainVoyager 20.6 (Brain Innovation, RRID:SCR_013057), Neuroelf v1.1 (www.neuroelf.net, RRID:SCR_014147), and in-house Matlab scripts (Mathworks, version 2018a, RRID:SCR_001622). Preprocessing of functional scans included slice timing correction (cubic spline interpolation), 3D motion correction by realignment to the first run image (trilinear detection and sinc interpolation), high-pass filtering (up to two cycles), smoothing (full width at half maximum (FWHM) = 2 mm), exclusion of voxels below intensity values of 100, and co-registration to the anatomical T1 images. Anatomical brain images were corrected for signal inhomogeneity and skull-stripped. All images were subsequently normalized to Montreal Neurological Institute (MNI) space (3×3×3mm functional resolution, trilinear interpolation).

### Estimation of cortical responses to each stimulus

A general linear model (GLM) analysis was applied. Each modeled predictor corresponded to one of the 24 individuals, and included all experimental trials where this individual’s name was shown, irrespective of the question asked in this trial. Predictors were convolved with a canonical hemodynamic response function and the model was fitted to the BOLD time-course at each voxel. 24 motion parameters were added to the GLM to eliminate motion-related noise; the six translation and rotation parameters, their temporal derivatives, and the squared values of the six parameters and their derivatives (Charest et al., 2018; Friston et al., 1996). The resulting beta values were converted to t-values using BrainVoyager contrasts (1 for each predictor and 0 for all other predictors) (Misaki et al., 2010). Finally, the t-values corresponding to each individual name were averaged across experimental runs to obtain one individual name-specific pattern (Dimsdale-Zucker and Ranganath, 2018).

### Definition of regions of interest (ROIs)

Functional ROI masks of spatial scene-selective regions (retrosplenial complex (RSC), parahippocampal place area (PPA) and occipital place area (OPA)) were obtained from a previous publication (Julian et al., 2012, http://web.mit.edu/bcs/nklab/GSS.shtml). These masks represent group activation clusters from 30 subjects who watched visual images with a contrast of scenes>objects. Functional ROIs for social and spatial proximity judgments were defined from the results of our previous study (Peer et al., 2015, contrasts - social judgments vs. lexical control and spatial judgments vs. lexical control, random-effects group analysis results). An anatomical hippocampal region-of-interest was extracted from the AAL atlas (Tzourio-Mazoyer et al., 2002).

### Representational similarity analysis (RSA)

Analyses were performed using CosmoMVPA (Oosterhof et al., 2016) and in-house Matlab scripts. The t-values for each of the 24 individuals were used as inputs for a spherical RSA searchlight with a radius of three voxels (Kriegeskorte et al., 2008). The mean of each voxel across all 24 conditions was subtracted from all voxel values (Diedrichsen and Kriegeskorte, 2017). For each searchlight sphere location, a neural dissimilarity matrix was computed between the activity vectors for the 24 individuals using Pearson’s correlation. The neural dissimilarity matrix was compared to each behavioral dissimilarity matrix (Facebook distance, personal affiliation, personality, and appearance dissimilarity matrices) using Spearman’s correlation (Nili et al., 2014). Group analysis was performed for each matrix’s correlation values using permutation testing (10,000 iterations) with threshold-free cluster enhancement, as implemented in the CosmoMVPA toolbox (Smith and Nichols, 2009; Stelzer et al., 2013).

To identify the independent contribution of each social distance matrix, a similar RSA searchlight was performed for each of the four dissimilarity matrices (social network distance and personal affiliation, personality and appearance dissimilarity matrices), after regressing out from each matrix the other three matrices (Parkinson et al., 2017). Group-level results were again computed using permutation testing with threshold-free cluster enhancement. This analysis was also performed in each of the ROIs for each subject (using the activity vectors across all ROI voxels), and significance in each ROI was computed using a one-sample one-tailed t-test (with FDR correction for the number of ROIs).

### Overlap with resting-state networks

Overlap was calculated between the significant voxels in the RSA searchlight group analysis result and each of the seven major resting-state networks as identified by Yeo et al., 2011 (Yeo et al., 2011, https://surfer.nmr.mgh.harvard.edu/fswiki/CorticalParcellation_Yeo2011). Overlap was calculated as the number of significant searchlight voxels included in each network divided by the overall number of significant searchlight voxels. Significance of overlap with each network separately was computed by permuting the voxel labels for the seven networks 1000 times and looking at the number of permutations reaching the same degree of overlap or higher.

### Visualization

Volume results were converted to surface representations and displayed using Connectome Workbench (Marcus et al., 2011).

### Data sharing

The full preprocessing codes, analysis codes and resulting statistical maps are available at https://github.com/CompuNeuroPsychiatryLabEinKerem/publications_data/tree/master/social_networks.

## Acknowledgments

This work was supported by the Israeli Science Foundation (Grant No. 1306/18). MP is supported by a Fulbright postdoctoral fellowship from the United States–Israel Educational Foundation, by a Zuckerman STEM Leadership Program fellowship, and by the Eva, Luis & Sergio Lamas Scholarship Fund. We wish to thank our study participants for their efforts; Assaf Yohalashet, Lee Ashkenazi, Dr. Yuval Porat and Prof. Leon Deouell from the ELSC neuroimaging unit for their help in MRI scanning; Catherine Nadar, Dr. Iva Brunec, Dr. Nicholas Diamond, Dr. Yoed Kenett and Dr. Zvi N Roth for helpful comments on the manuscript; and Prof. Russell Epstein (University of Pennsylvania) for helpful suggestions and discussions.

## Author contributions

Conceptualization, MP and SA; Methodology, MP and SA; Investigation and formal analysis, MP, MH, BT and SA; Visualization, MP and MH; Writing, MP and SA; Supervision and Funding Acquisition, SA.

## Supplementary materials

**Figure S1:**
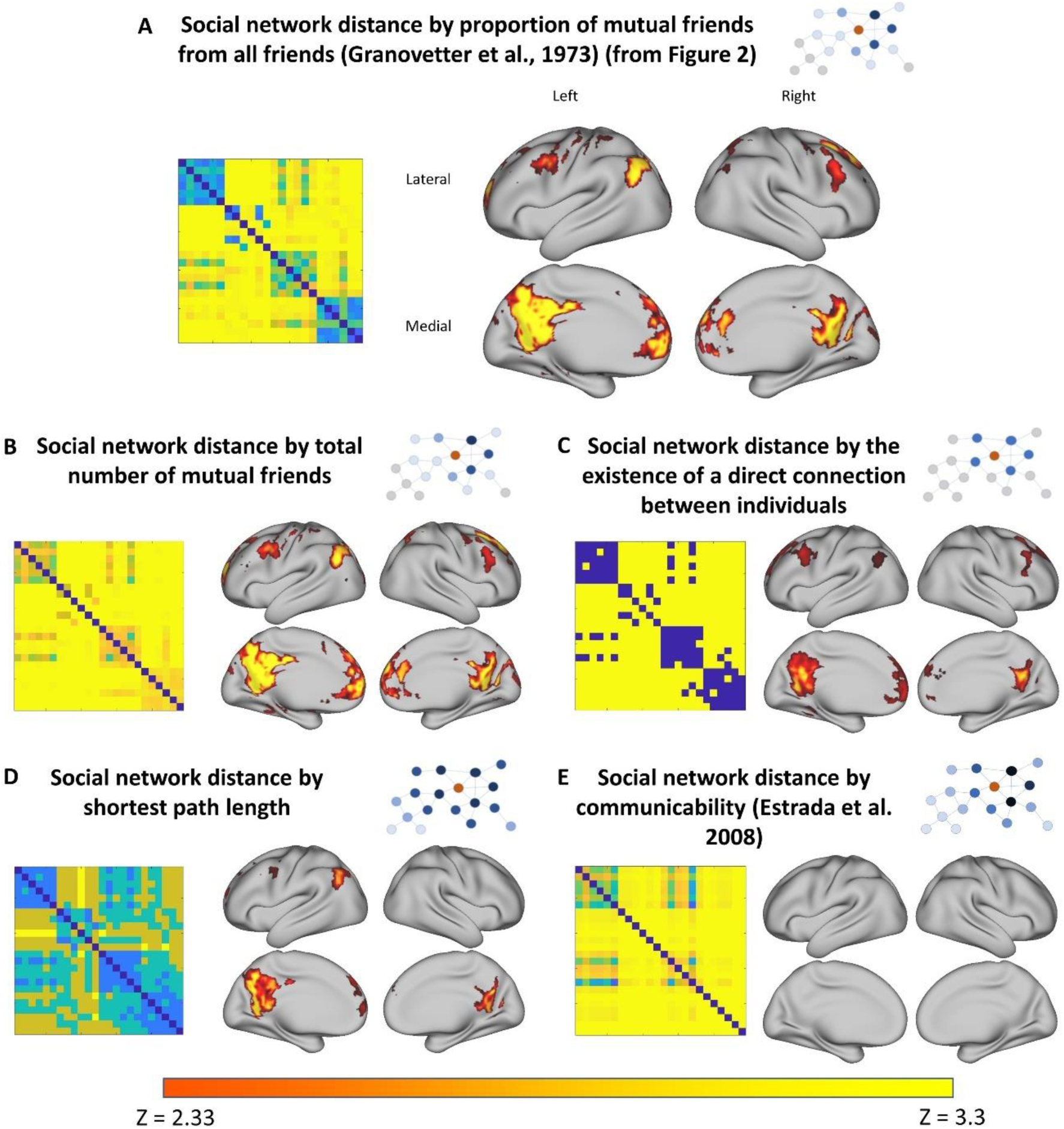
information on social network structure obtained using different social distance measures. Representational similarity searchlight on social media network connections between individuals. A) Proportion of mutual friends (similar to Figure 2), B) total number of mutual friends, C) direct connectivity between individuals (binary coding), D) shortest network path length, E) graph communicability (Estrada et al. 2008). Voxels with significant representational similarity are displayed for each social distance measure (p<0.01, Monte-Carlo permutation test, threshold-free cluster enhancement correction for multiple comparisons). Social distance matrices for each measure, from one sample subject, are presented to the left of each fMRI result (all matrices normalized to the range 0-1). Results are similar between different measures of social distance, except for the communicability measure which does not show any significant coding across the brain, indicating that pattern similarity reflects proximity between individuals in the social network instead of ease of information transfer across the network.

**Figure S2:**
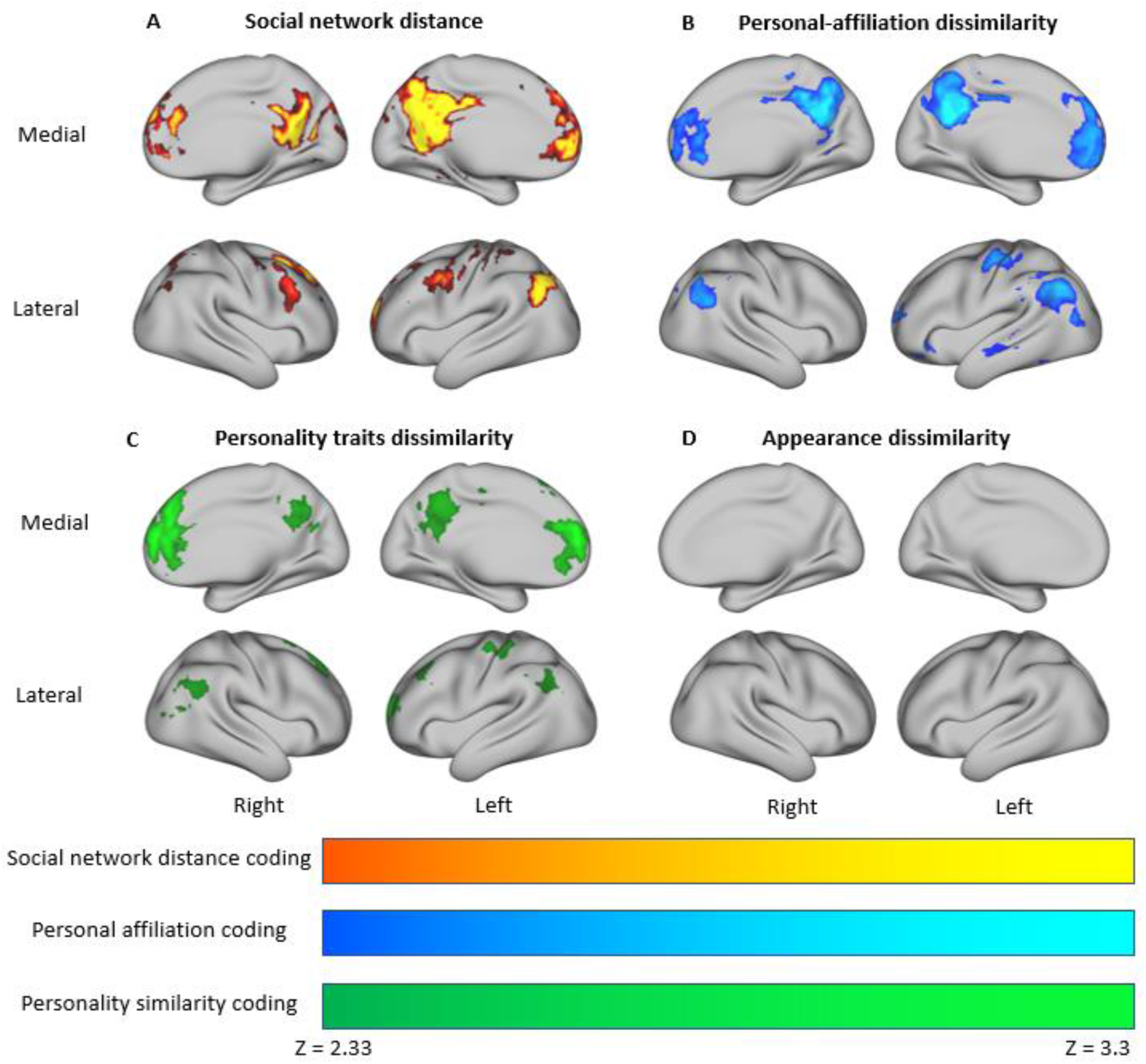
the default-mode network represents information on similarity between personally-familiar people across multiple aspects of social knowledge. Similarity of activity patterns related to thinking about familiar individuals was compared to the similarity between these individuals in different social aspects using a representational similarity searchlight analysis (sphere radius = 3 voxels). Pattern similarity across regions of the default-mode network was associated with: A) proximity between individuals in the subjects’ personal social networks, as derived from social media (Facebook) connectivity data (similar to Figure 2); B) similar ratings of personal affiliation to the subject; and C) similar ratings of personality traits; but not with D) similar ratings of appearance. All p-values<0.01, Monte-Carlo permutation test, threshold-free cluster enhancement correction for multiple comparisons.

**Table S1:** average within-subject correlation between the different dissimilarity matrices representing different aspects of social knowledge. The distance between individuals in the social network was found to be correlated to the similarity of responses to questions about personal affiliation, personality traits, and appearance. All correlations between dissimilarity matrices are significantly larger than zero except between the social network distance and the responses to appearance questions (all p<0.05, one-sample one-tailed t-test across subjects, FDR-corrected for multiple comparisons).

|  | Social network proximity (proportion of mutual friends) | Similarity in personal affiliation ratings | Similarity in personality ratings | Similarity in appearance ratings |
| --- | --- | --- | --- | --- |
| Social network proximity (proportion of mutual friends) | 1.00 | 0.13 | 0.08 | 0.03 |
| Similarity in personal affiliation ratings | 0.13 | 1.00 | 0.18 | 0.08 |
| Similarity in personality ratings | 0.08 | 0.18 | 1.00 | 0.08 |
| Similarity in appearance ratings | 0.03 | 0.08 | 0.08 | 1.00 |

**Figure S3:**
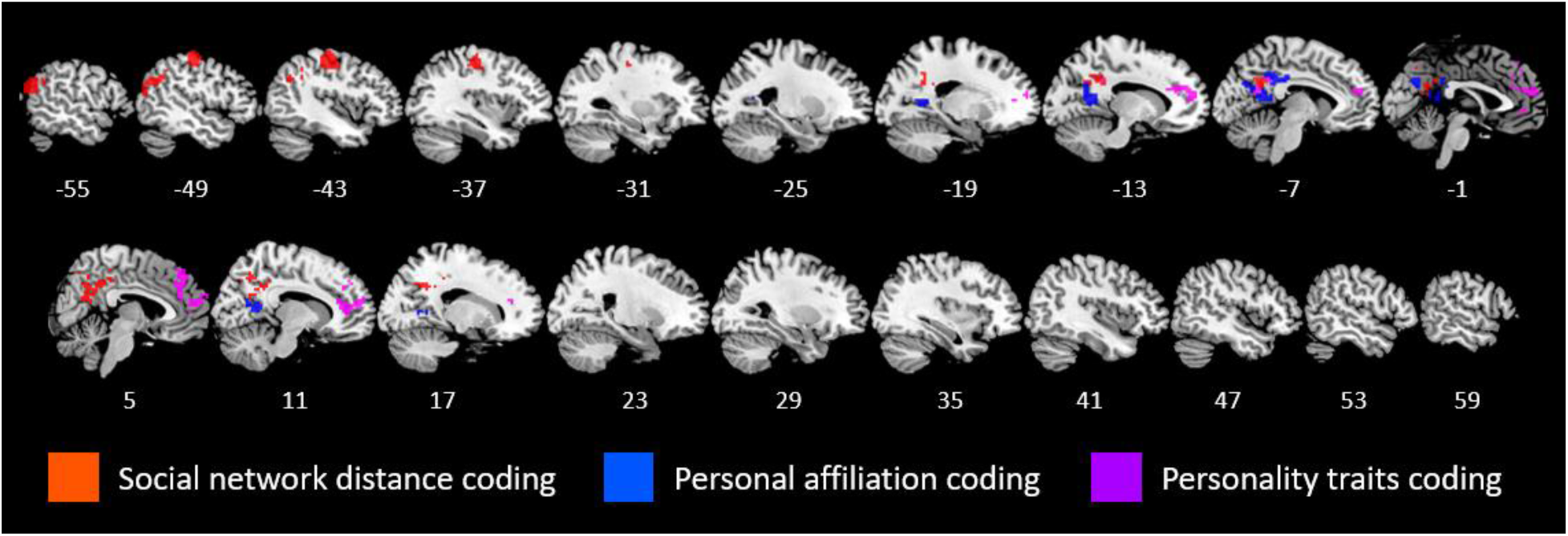
Coding of distinct aspects of social knowledge – multi-slice view. The similarity matrices for individuals’ social network distance, rated personal affiliation to the subject, rated personality traits, and rated appearance were regressed from one another and then used in a representational similarity searchlight, to identify the independent variance explained by each. Representational similarity searchlight, spherical radius = 3 voxels, p<0.01, Monte-Carlo permutation test, threshold-free cluster enhancement correction for multiple comparisons.

